## Supplementary material for "Unraveling the Role of Ctla-4 in Intestinal Immune Homeostasis: Insights from a novel Zebrafish Model of Inflammatory Bowel Disease": Animal Welfare and Ethics

浙江大学 ZHEJIANG UNIVERSITY

浙江大学实验动物福利伦理审查委员会
Laboratory Animal Welfare and Ethics Review Committee, ZJU

### 浙江大学实验动物福利伦理审查同意书

Approval of Animal Use Protocol,IACUC,ZJU 有效期：4年

| 项目批准号 AP CODE:ZJU20240828 | | 伦理申请编号 Appl No:30542 | |
|---|---|---|---|
| 实验名称<br>Protocol Title | 斑马鱼Ctla-4对肠道免疫稳态维系及其与炎症性肠病的相关性研究<br>Role of Zebrafish Ctla-4 in Maintaining Intestinal Immune Homeostasis and Its Relevance to Inflammatory Bowel Disease | | |
| 院系（部门）<br>Department | 浙江大学生命科学学院<br>College of Life Sciences, Zhejiang University | 合作单位<br>Co-organisation | no |
| 实验负责人<br>Principal Investigator | 邵建忠 | 申请人<br>Applicant | 秦露露 |
| 实验参与者<br>Participant(s) | 秦露露 | 动物信息<br>Animals to be used | 斑马鱼 (AB系、56) |
| 计划执行时间<br>Period of Protocol | 2024-10-08 -- 2024-12-05 | 实验场地<br>Animal Experiment Site | 浙江大学疾病模拟与模式动物平台 |
| 投票结果<br>Voting Results | 参会 Attendance : 6<br>不同意 Disagreed: 0<br>* 是否启动紧急程序 If the urgency process used: ☐ 是 Yes ☑ 否 No | 同意 Agreed: 5<br>弃权 Waiver : 0 | |
| 审查结果<br>Result of Review | ☑ 批准：符合动物福利伦理要求，允许开展实验 Approved<br>☐ 附条件批准：经调整方案后允许开展实验 Conditionally Approved | | |

评审意见 Comments:

该动物实验方案经过浙江大学实验动物福利伦理审查委员会审核，符合动物福利和伦理原则，符合国家实验动物福利伦理的相关规定，允以批准实施。项目负责人须依据承诺和既定实验方案开展科学研究，确保实验动物福利伦理原则的真正实施。 This animal use protocol listed below has been reviewed and approved by Laboratory Animal Welfare and Ethics Review Committee of Zhejiang University, and is following the principles of animal welfare and ethics, as well as relevant national regulations. The protocol should be performed in accordance with the approved protocol; the researchers have committed to guarantee the animal welfare during the whole procedures.

主任委员 Committee Chair:

日期 Date: 2024/10/24

签发单位 Issuing unit: 浙江大学实验动物福利伦理审查委员会

Laboratory Animal Welfare and Ethics Committee of Zhejiang University
